# Validation of a high-throughput screening assay for the characterization of cryoprotective agent toxicity

**DOI:** 10.1101/2025.05.26.654916

**Authors:** Justyna J. Jaskiewicz, Abby Callahan-Muller, Nora Gaby-Biegel, Zachary Glover, Rebecca D. Sandlin

## Abstract

Cryoprotective agent (CPA) toxicity is a major limitation in cryobiology. The discovery of new CPAs and formulation of low toxicity CPA cocktails can be streamlined by use of high-throughput screening (HTS) assays. HTS allows for rapid examination of a large chemical space, but assays should be rigorously validated to ensure the production of high-quality datasets. Here, we report the development of a CPA toxicity assay, validation of the assay for HTS using prevailing statistical metrics, and completion of a pilot screen to assess assay performance against various CPA cocktail formulations. The resulting assay, which consists of T24 cell monolayers in 96-well microtiter plates, was found to exhibit favorable performance including a Z-factor of 0.75, with intra-assay coefficient of variation and drift of <20%, which indicates that the assay is suitable for HTS implementation. A pilot screen was then completed using 587 unique CPA cocktails with concentrations ranging from 3.5-8 M. Each cocktail was tested with multiple replicates to characterize plate-to-plate and day-to-day variability of the assay (N=2-9, 2,352 total experiments). Of the 56 plates examined in the pilot screen, 53 exhibited favorable performance based on assessment of the on-plate controls (95% success rate). CPA cocktails tested at a concentration of 5 M were found to be highly informative, where cell survival ranged from 2.8-87.3%, with a favorable hit rate of 1.7%, defined here as cell viability ≥80%.

## 1. Introduction

High-throughput screening (HTS) assays are developed with the intention of rapidly screening large numbers of compounds in microtiter plate format. The goals of HTS include: to achieve rapid and efficient screening of thousands to millions of unique compounds; to generate large datasets for the purpose of identifying patterns and relationships between chemicals and their biological activities; the identification of “hits” (i.e. active compounds); and to explore a diverse chemical space. The resulting dataset can then be used for lead identification and optimization of the hits or to help understand mechanisms of action. However, the ‘garbage-in, garbage-out’ concept applies [4], where poorly designed HTS assays may lack sensitivity or reproducibility leading to the production of unreliable datasets. To ensure the developed assay is high quality, validation studies are typically completed that incorporate analytic quality control methods to assess assay quality. [11] This includes the identification of appropriate positive and negative on-plate controls, establishment of plate uniformity and reproducibility, and quantification of plate drift. Further, statistical methods are utilized to assess assay quality including the Z-factor [23] and coefficient of variation (CV). The Z-factor measures the statistical effect size of positive and negative controls, where a Z-factor between 0.5-1 is considered an excellent assay due to the ability to reliably distinguish hits in large datasets based on their activity relative to the positive and negative on-plate controls. CV and drift are also applied to understand variability among controls, where a CV and drift <20% is desired. Collectively, these metrics can validate assay quality and ensure reproducibility for the collection of high quality and informative data. This validation is further expected to reduce the false-positive and false-negative hit rates by ensuring minimal plate-to-plate or well-to-well variability. Following successful assay validation, a pilot study (also called a mini-validation study), is typically performed prior to the primary screen to demonstrate assay logistics. [5] This process consists of screening a smaller library to identify barriers to assay implementation such as high hit rates or poor reproducibility. [5,17] Continued quality control during the pilot screen is accomplished using on-plate positive and negative controls to serve as plate acceptance criteria using the Z-factor method. Here, only plates with a Z-factor of >0.5 are considered acceptable. Assuming the pilot screen demonstrates robust performance, the much larger primary screen is then completed which includes thousands to millions of test wells. Hits from the primary screen are then selectively chosen to confirm activity before proceeding to secondary screening. This process, known as cherry-picking [16], involves selecting most promising candidates from a larger set for further validation and optimization, ensuring that only the most promising compounds advance in the discovery process. The resulting dataset is then used to select hits for lead optimization in the ‘hit-to-lead’ process. Schematic 1 overviews this typical HTS process. [5,18]

**Schematic 1.**
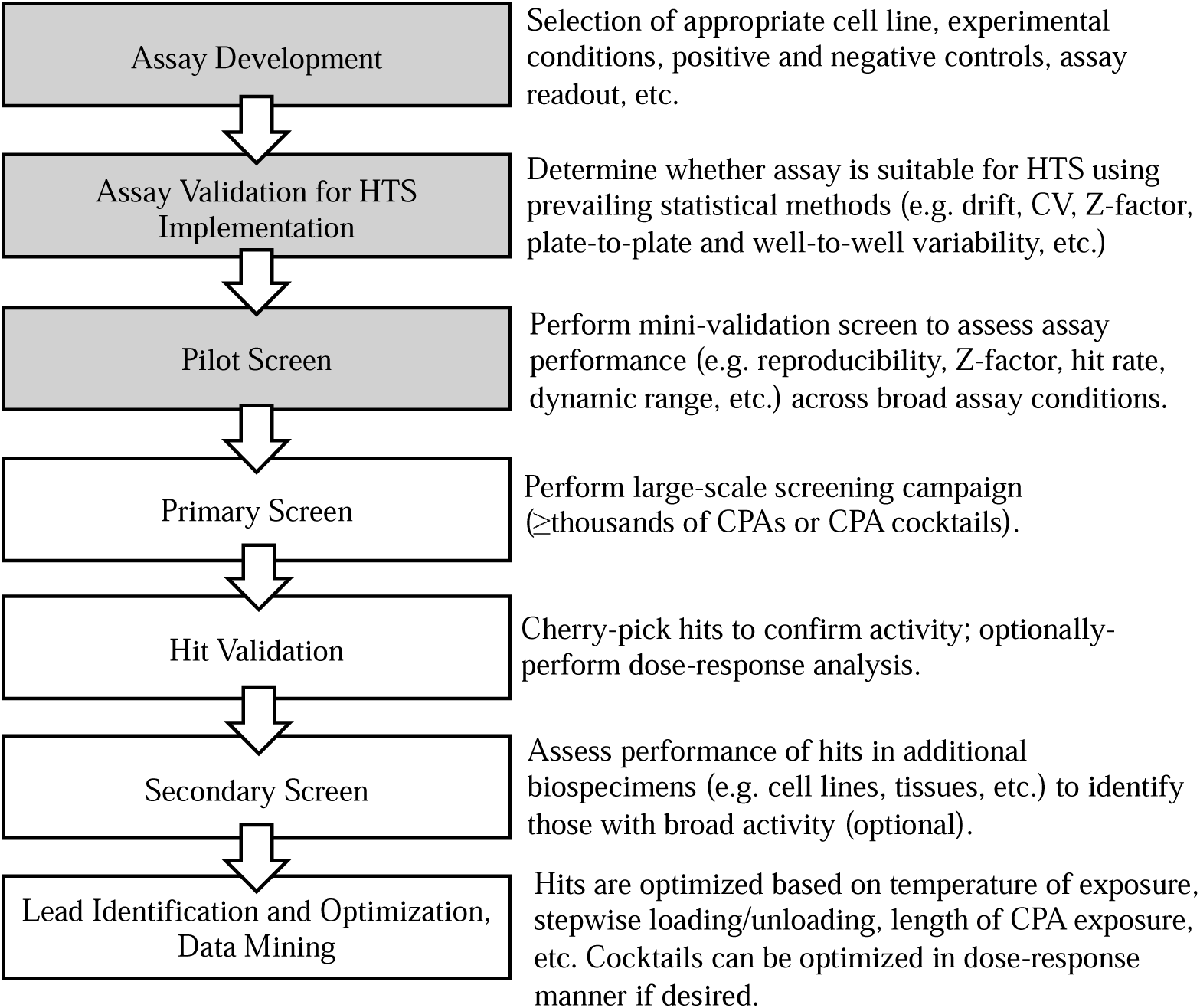
**Flow chart of typical HTS process for lead discovery**. The first column reflects typical approach to HTS used in the drug discovery field, while the second column overviews how this process can be applied in the context of CPA or CPA cocktail discovery. Highlighted boxes represent the progress reported in this manuscript.

HTS assays could be utilized to facilitate the rapid identification of novel cryoprotective agents (CPAs), where multicomponent CPAs are of particular interest as cocktails exhibit reduced toxicity compared to single component CPAs at the same overall concentration [2,8]. Intracellular CPAs (e.g. glycerol, ethylene glycol, etc.) are molecules that penetrate cell membranes and protect biospecimens from damage during cryogenic storage yet are typically toxic at concentrations required for cryopreservation. This is especially true for vitrification, an ‘ice-free’ approach to cryopreservation where relatively high concentrations of CPAs are loaded into biospecimens followed by rapid cooling through the glass transition, resulting in amorphous solidification of water (i.e. glass formation). Recently, several high-throughput assays have been adapted to 96-well microtiter plate to more efficiently quantify CPA and CPA cocktail toxicity using cell monolayers [10,21,22], non-human organs [13], or whole organisms [7]. These assays offer a promising approach towards scale-up of toxicity screening and have been used to examine from 24-340 unique CPA conditions. While these assays show great promise towards the use of HTS for CPA optimization and discovery, it remains unclear whether a CPA toxicity assay can be developed to meet the rigorous validation metrics applied in the HTS field.

To our knowledge, no prior CPA toxicity assay has been reported which adheres to the prevailing standards for HTS assay development and validation for implementation. Given the increased interest in rapid screening for CPA toxicity, the goal of this study was to develop an assay to screen for CPA toxicity, perform an HTS validation study, and complete a pilot screen – the first three steps in the typical HTS discovery process (Schematic 1). The validation stage is especially critical, as the actual scaling of assays from a few dozen or hundred CPA solutions to thousands or millions of CPA solutions demands rigor to ensure the data collected is informative and reproducible. The developed assay, which consists of a monolayer of T24 cells in 96-well format, was found to perform favorably for HTS implementation based on Z-factor, CV and drift as measured during assay validation stages. The pilot screen then included 587 unique CPA cocktails containing between 2-7 individual CPAs. To characterize assay variability, replicates were used (N=2-9), resulting in efficient execution of 2,352 total experiments.

## 2. Materials Methods

### 2.1 Cell culture and cell lines used

All cells were maintained in monolayer culture in a 5% CO_2_, 37 °C incubator in cell-specific culture medium supplemented with 10% FBS with penicillin (100 U/ml) and streptomycin (100 µg/ml). Cells were passaged at a ratio and frequency instructed by the manufacturer unless specifically indicated. For passaging, cell monolayers were detached by treatment with trypsin (0.05%, 7-15 min), washed with respective media and centrifuged (199xg, 5 min).

HEPA 1-6 mouse hepatocellular carcinoma cells (ATCC, cat. No. CRL-1830) and MSC bone marrow-derived human mesenchymal stem cells (ATCC, cat. No. PCS-500-012) maintained for up to five passages were grown in DMEM medium; HCT-8 human colorectal adenocarcinoma cells (ATCC, cat No., CCL-244) in MEM medium, LnCaP human prostate carcinoma (ATCC, cat. No. CRL-1740), PC3 human prostate adenocarcinoma (ATCC, cat. No. CRL-1435), NCI-H1650 human bronchoalveolar carcinoma (ATCC, cat No. CRL-5883) and MDBK bovine kidney epithelium (ATCC, cat. No. CCL-22) cells in RPMI medium; and SK-BR-3 human breast adenocarcinoma cells (ATCC, cat. No. HTB-30) in McCoy’s 5A medium. HUVEC human primary umbilical vein endothelial cells (ATCC, cat. No. PCS- 100-013) were grown in F-12K media (ATCC, cat. No. 30-2004) supplemented with endothelial cell growth supplement (Fisher Scientific, cat. No. CB-40006) and heparin ( 100 µg/ml) and were maintained for up to ten passages. The T24 human urinary bladder cell line (ATCC, cat. No. HTB-4) and T24-GFP variant stably expressing GFP transgene (Applied Biological Materials, Cat No. T3912) were grown in McCoy’s 5A medium (ATCC, cat. No. 30-2007). Both T24 cultures were passaged 3 times a week at a 1:4-1:7 ratio and were maintained for up to 25 passages while monitored for consistent doubling time (∼20h).

### 2.2. Identification of HTS suitable cell line

Adherent cell lines were seeded on clear culture-treated 96-well plates (Corning) at a density leading to confluency on the day of CPA challenge. Media was removed from wells by flicking the plate, followed by dabbing against paper tissue to ensure removal of media residue. To avoid desiccation of monolayers CPA cocktails prepared in PBS-10% FBS was added immediately in a single step and was incubated for 60 min at ambient temperature (22-25°C). CPA was then removed in 3 steps; first by 2-fold dilution with PBS followed by 1 min incubation, additional 4-fold dilution with PBS followed by 2 min incubation and then replacement with respective cell media. Monolayers were imaged before CPA challenge, 30 min after CPA challenge and immediately following CPA dilution.

### 2.3 Automated liquid handling

All steps requiring deposition of the liquid into a 96-well plate, with exception of the transfer of CPAs from the source plate to the experimental plate, were performed using a MultiFlo FX multi-mode dispensing robot (Biotek Instruments, Inc., Winooski, VT, USA). The preparation of multi-component CPAs, along with cell seeding, was completed using a single tip dispensing RAD cassette with a 5 µl volume increment (Agilent Technologies, cat. No. 1260016). CPA dilution and addition of MTT assay reagents were performed using an 8-tip dispensing cassette with a 1 µl volume increments (Agilent Technologies, cat. No. 7170012).

### 2.3 Overview of high throughput CPA screening assay

The screening assay is conducted over a period of three days, encompassing three key steps as illustrated in Figure 3A: cell preparation, CPA challenge, and viability assessment. The CPA dose-response toxicity assay shown in Figure 4 was performed following the same procedural steps.

**Figure 1.**
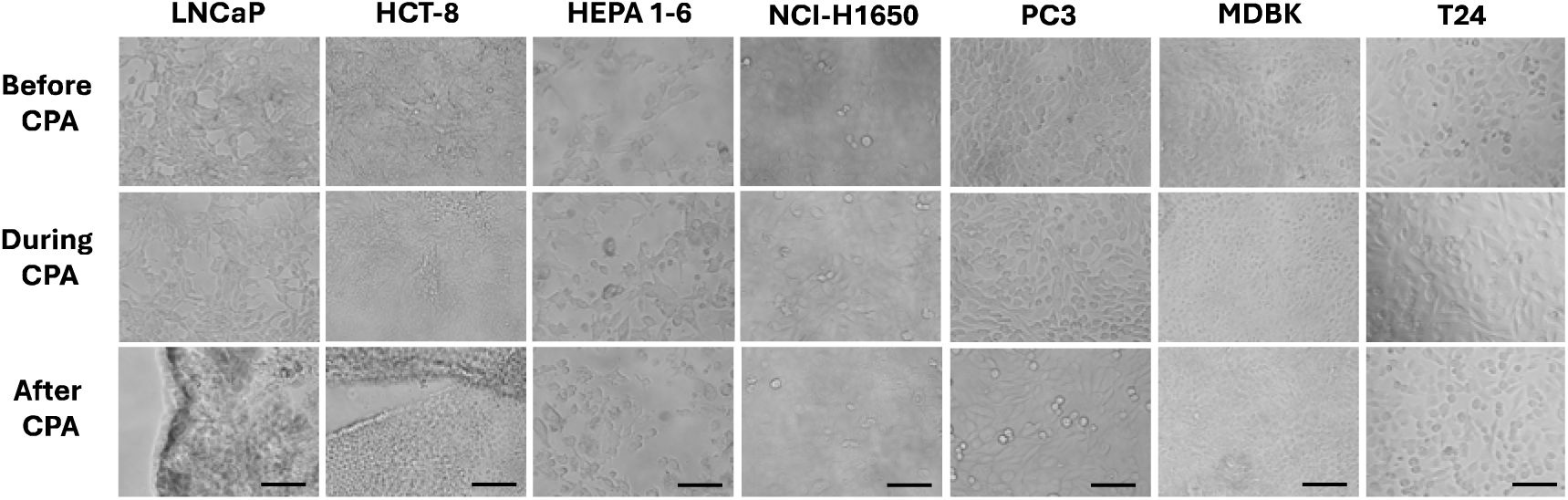
**Selection of appropriate cell lines for the CPA toxicity HTS assay**. Ten cell lines were selected to assess for compatibility in the CPA toxicity assay as summarized in Supplemental Table 1 (seven are shown here). LNCaP and HCT-8 exhibit significant cell detachment upon CPA dilution, while HEPA 1-6 fails to reach confluence, excluding them from consideration in this study. NCI-H1650, PC3, MDBK, and T24 all tolerate single-step CPA addition without observed detachment. However, NCI- H1650 is excluded due to slow replication rate, PC3 due to incompatibility with MTT assay development, and MDBK due to sensitivity to 1,2-propanediol. Consequently, T24 was selected for development of the HTS CPA toxicity assay. Scale is 100 µm.

**Figure 2.**
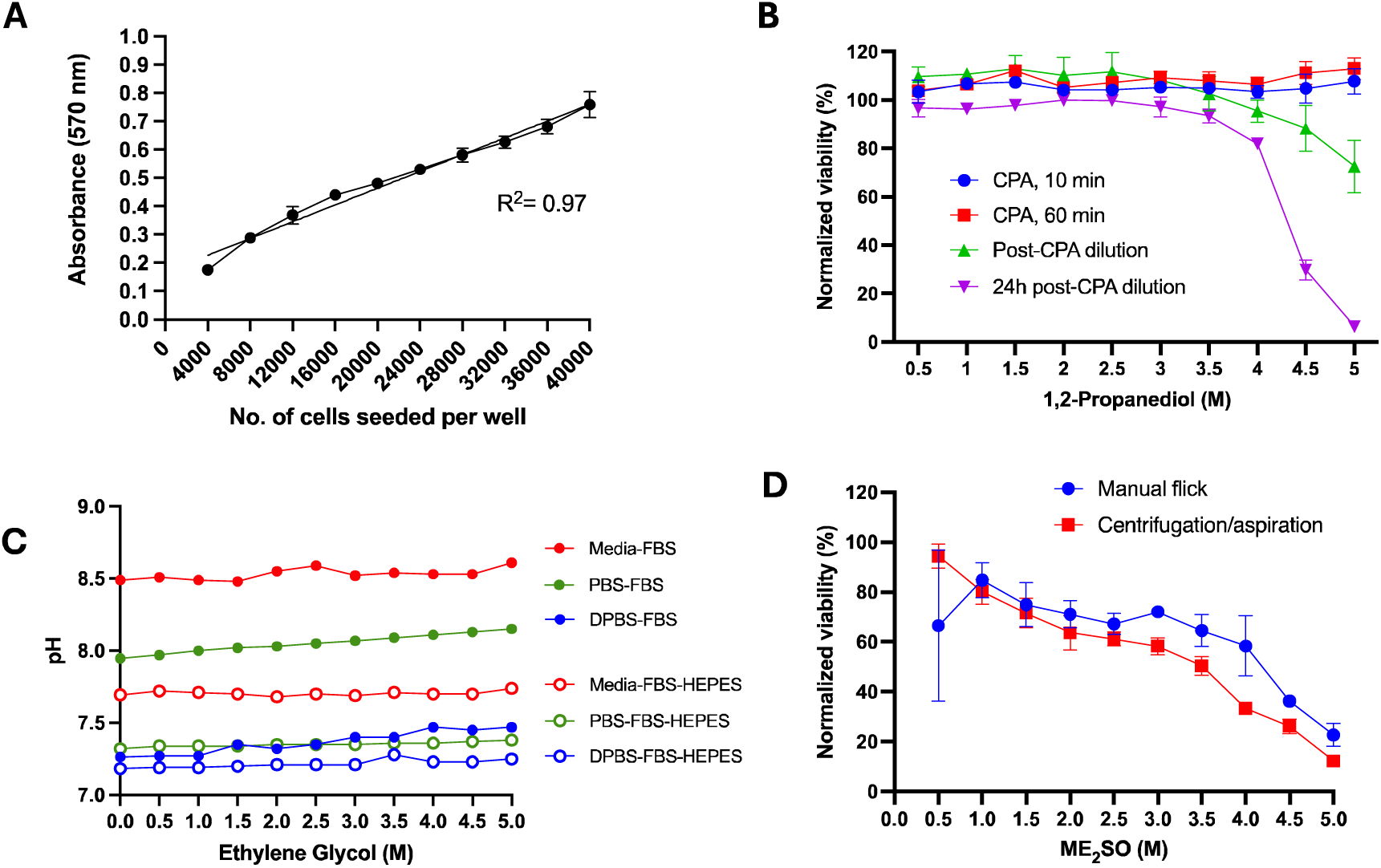
Validation of assay parameters. (A) The T24 standard curve was measured using the MTT viability assay. T24 cells were seeded serially at densities ranging from 4,000-40,000 cells/well. These densities are within the linear range of the MTT viability assay (R^2^=0.97). (B) The kinetics of CPA toxicity was monitored in the T24-GFP cell line using 1,2-propanediol. The late onset of toxicity is observed 24 h after CPA dilution. (C) Multiple vehicle carrier solutions were evaluated to assess the effects of CPAs on pH, where ethylene glycol is shown (additional CPAs in Supplemental Figure 4). (D) A comparison of a manual and automated method for CPA removal from the plates was performed. The automated aspiration of diluted CPAs after centrifugation delivered more consistent results compared to the manual method.

**Figure 3.**
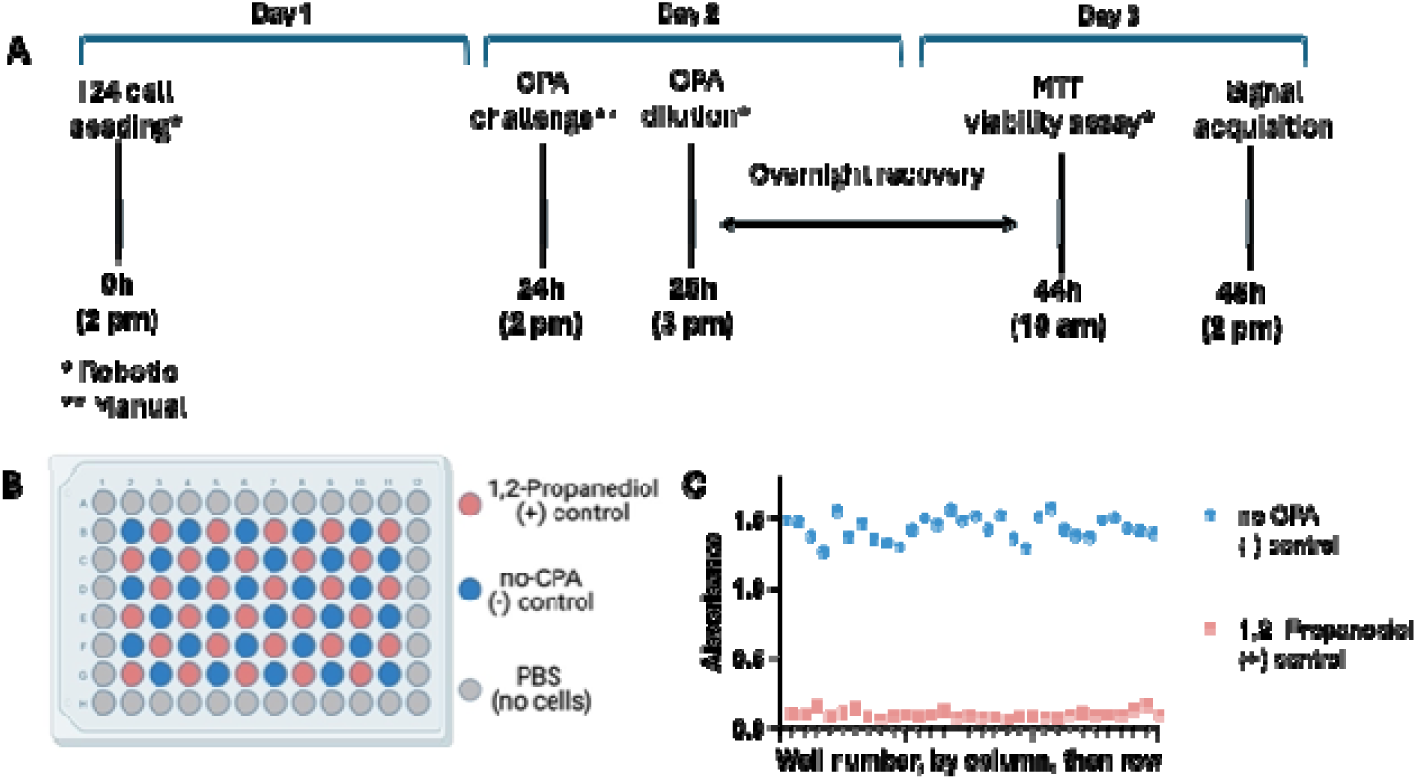
Validation of the CPA toxicity HTS assay. **(A)** The HTS CPA toxicity assay consists of three key steps: T24 cell preparation (day 1), CPA challenge (day 2) and assessment of viability (day 3). **(B)** A checkerboard plate layout was used for evaluation of HTS assay performance, which consisted of alternating control wells containing a ‘low’ assay signal (5 M 1,2-propanediol, positive control) and ‘high’ assay signal (no-CPA, negative control). Figure created with BioRender.com. **(C)** A comparison of the high and low assay signal was plotted for assessment of assay quality and reliability. The maximal drift of the positive control was 7.5% and the negative control was 10.3%. An intra-assay coefficient of variation (CV) was calculated at 5.9% for the negative and 22.5% for the positive control. Standard deviation (SD) of the low signal control is 3 times less than that of the high signal control, meeting the criteria for plate acceptability.

**Figure 4.**
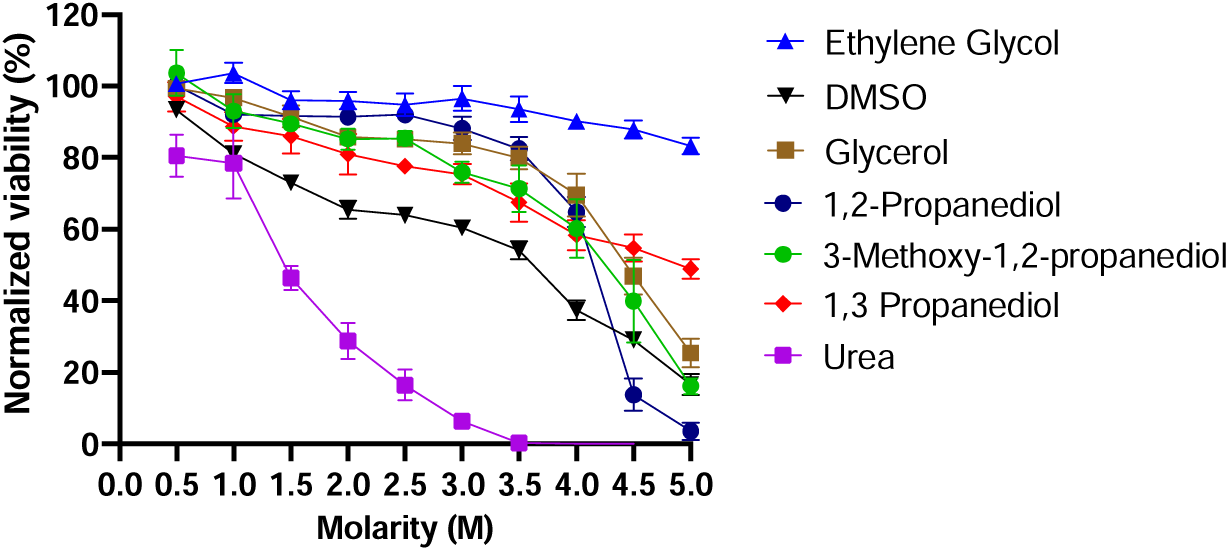
Dose response toxicity of CPAs selected for this study. To quantify the individual toxicity effects of each CPA, the dose-response toxicity was assessed in the T24 CPA toxicity assay. Each CPA was tested from a range of 0.5-5 M in 0.5 M increments.

#### Day 1

*Cell seeding.* Trypsinized T24 cells were suspended in a Falcon tube and seeded on a clear culture-treated 96-well plate (Corning) at 20,000 or 25,000 cells per well using a liquid dispensing robot and a single tip dispensing cassette. If T24-GFP were used, they were seeded at on black bottom black culture-treated 96- well plates (Greiner) at 25,000 cells per well. Cells were dispensed into each experimental well at a volume of 100 µl. The wells on the perimeter of the plate were filled with 100 µl of PBS to account for possible edge effects. Seeded plates were incubated overnight until confluency was reached at an estimated density 40,000 cells/well.

#### Day 2

*CPA source plate preparation.* CPA cocktails were prepared in a carrier solution consisting of DPBS supplemented with 10% FBS and 25 mM HEPES, unless otherwise indicated. A source plate with various CPA cocktails was prepared in a 96-well plate freshly prior to the cell challenge. The components of CPA cocktails were dispensed into wells using a single tip dispensing RAD cassette. The layout of a source plate was designed as shown in Figure 5A, such that positive and negative control wells were uniformly distributed across the plate allowing for downstream assessment of assay quality. Following exclusion of edge wells and incorporation of control wells, 42 wells remain available on a plate for screening of CPA cocktails. After dispensing, the plate was sealed with a plastic film and agitated on a shaking rotor for 20 min at 100 rpm.

**Figure 5.**
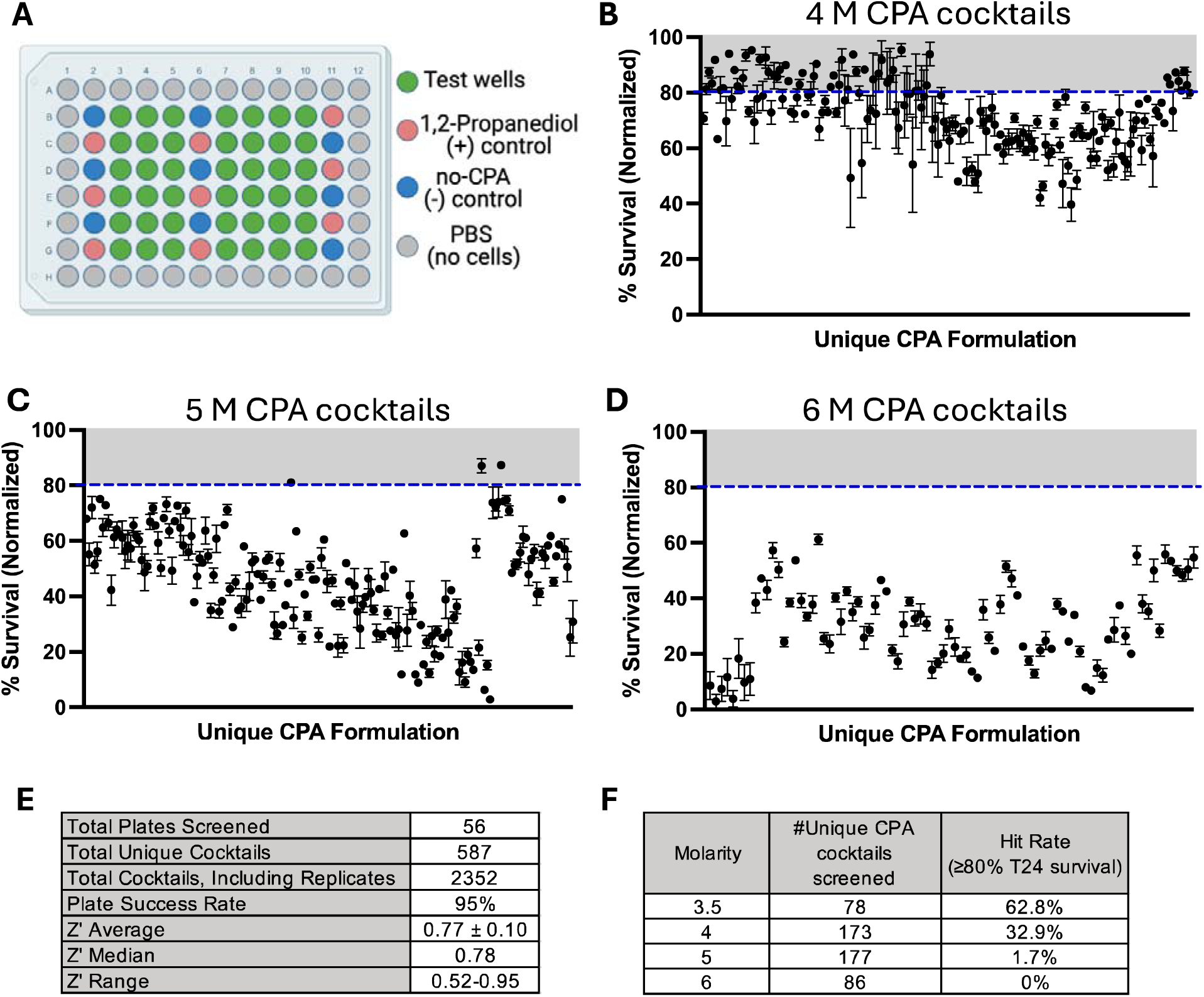
Results of the pilot screen to quantify toxicity of CPA cocktails. **(A)** The plate layout for CPA toxicity HTS assay in 96-well format is shown, which consists of 42 experimental and three columns of alternating positive and negative control wells. Figure created with BioRender.com. **(B, C, D)** The pilot screening results of unique CPA cocktails tested at 4, 5 or 6 M concentrations are shown, where each cocktail contained between two and seven individual CPA components. Observed clustering or patterns in cell survival are due to similarities in the composition of neighboring CPA cocktails. See supplemental file for detailed list of CPA cocktail compositions and associated viability values. **(E)** An overview of key results from the pilot screen are summarized. **(F)** The resulting hit rates for CPA cocktails tested at each indicated overall concentration are shown, where hits are defined as CPA cocktails that lead to ≥80% T24 survival.

*CPA challenge.* Both the CPA source plate and T24 experimental plate were equilibrated at room temperature for 30 min. Media was removed from wells by flicking the plate, followed by dabbing against paper tissue to ensure removal of media residue. The following steps were performed promptly to prevent desiccation of monolayers. In a single-step CPA challenge protocol, 50 µl of 1x CPA cocktails were dispensed by transferring them from the source plate using a multi-channel pipette. In the multi-step CPA challenge protocol, the initial step involved automated dispensing of 25 µl of the carrier solution using an 8-tip dispensing cassette. Following was the incremental deposition of 25 µl 2x CPA cocktails by transferring them from the source plate using a multi-channel pipette, resulting in a cumulative volume of 50 µl in each well. Following deposition of CPAs, the edge wells were filled with 50 µl PBS and the plate was left at room temperature to incubate for 1h. The selected conditions led to an appropriate dynamic range for selecting hits among CPA cocktails examined at 5 M (Fig. 5C).

*CPA dilution and removal.* Following the challenge, the CPA underwent a three-step dilution protocol using automated dispense and aspiration robots. First, 50 µl of the carrier solution was added with an 8-tip dispensing cassette. After a 1 min incubation, an additional 100 µl of carrier solution was added and allowed to incubate for 2 min. The plate was then centrifuged at 50 x g for 1 min and contents were aspirated using an automated plate aspirator (BioTek ELx50). The nozzle was set to hover 3 mm above the monolayer and aspirate at the speed 5 mm/sec. Lastly, wells were refilled with 100 µl of complete cell media and plates were incubated overnight at 5% CO_2_ at 37 °C.

#### Day 3

*Viability assay.* Cell survival was typically measured by means of the colorimetric 3-(4,5-dimethylthiazol- 2-yl)-2,5-diphenyltetrazolium bromide (MTT) reduction assay. In some cases, viability was assessed by monitoring the fluorescent signal of the T24-GFP cell line. All steps requiring deposition of the liquid were performed using a multi-mode dispensing robot.

The MTT assay was performed as described elsewhere, with modifications. [20] Briefly, cell media was removed from wells by flicking the plate. A solution of MTT reagent suspended in phenol-free and FBS-free media at a concentration of 5 mg/ml was added to the plate and incubated for 3 h at 37 °C. After removal of the MTT reagent by gently flicking the plate, cells were solubilized in a solution consisting of 2-propanol/0.3 mM hydrochloric acid /0.5% sodium dodecyl sulfate for 30 min while agitating on an orbital shaker followed by 10 min on a platform rocker. Viability was calculated from the absorbance measured at 570 nm using a microplate reader (BioTek), where % viability was then calculated relative to the on-plate negative (no-CPA) controls.

For measurement of the T24-GFP viability, cell media was replaced with PBS and the fluorescence signal was acquired using a microplate reader (excitation 485/20, emission 520/20, gain 120, BioTek). The signal was adjusted to background and normalized to the positive control wells corresponding to 100% viability.

Viability of cells in suspension was assessed using a live/dead stain and detected through fluorescent microscopy. Briefly, cells were stained with 12.5 µM Calcein AM and 0.1 µg/ml propidium iodide for 5 min in the dark. Subsequently, cells were examined and enumerated under 200x magnification using an EVOS microscope.

### 2.4. Statistical characterization of assay performance

For the purpose of assay validation, positive and negative controls were added to the plate in an alternating, checkerboard pattern. Assay quality was then assessed by calculation of the Z-factor, [23] drift and CV as described in the guidelines proposed by the National Institutes of Health Chemical Genomics Center assay guidance manual. [4]

The %CV for the control plate was calculated for the both positive and negative controls as follows:

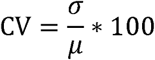

where σ is the standard deviation of the positive or negative controls, and µ is the average signal of the positive or negative controls.

The Z-factor (Z’) was calculated as

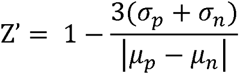

where (J_p_ and (J_n_ are the respective standard deviations of the positive and negative controls, and µ_p_ and µ_n_ are the means of the positive and negative controls, respectively. A Z-factor was calculated for each experimental plate using the three columns of positive and negative controls on each plate, which served as a quality control for plate acceptance criteria in the pilot screen.

The percent signal drift was calculated both by row and by column. For example, the column drift was calculated as

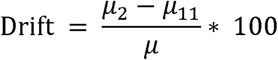

where µ_2_ and µ_11_ are the average signals of the outermost columns 2 and 11, respectively, and µ is the average of these two columns.

### 2.5 Dose-response toxicity and CPA pilot screen

Using the assay described above, a dose-response curve was evaluated to establish the contributing toxicity of individual CPAs. Next, a library consisting of 587 unique CPA cocktails at an overall combined concentration of 3.5-8 M was constructed. Each cocktail consisted of at least two out of seven components, namely ethylene glycol, Me_2_SO, glycerol, 1,2- propanediol, 3-methoxy-1,2-propanediol, 1,3-propanediol, and urea. The library was constructed to include a wide range of CPA combinations and concentrations and to characterize the overall survival of cells at a range of concentrations (see Supplemental Excel file).

## 3. Results and Discussion

### 3.1 Assay Development: Identification of suitable cell line for CPA toxicity HTS assay

Typically, in HTS, molecules are screened at concentrations in the nM-µM range. To examine CPA toxicity, however, we are interested in molar concentrations. This could create a challenge as high concentrations of CPAs are expected to induce an osmotic ‘shrink-swell’ response [14]. Similarly, dilution of CPAs would lead to a ‘swell-shrink’ response. Thus, if CPA loading and unloading were to occur too rapidly, cell volume changes could compromise both viability and attachment to the plate surface. This creates a bottleneck for HTS design, as the assay should be simple to execute, and multi-step loading/unloading would significantly reduce throughput. While the incorporation of well-to-well transfer robots (e.g. Myra, Tecan, etc.) can deliver CPAs in multiple steps, this can be cumbersome when the concentration of CPA and the number of plates screened on a given day is high. For example, examination of a 96-well microtiter plate at a concentration of 5 M CPA would require five stepwise additions to maintain the osmotic gradient at ∼1,000 mOsmol. While this is practical when using a small number of plates, it becomes logistically challenging when the assay format is scaled for the primary HTS screen (Schematic 1) and the goal is to execute many thousands of experiments. Selection of a cell line that tolerates high concentrations of CPAs in a single step without exhibiting cell detachment or excessive mortality was therefore central to development of the assay to ensure downstream HTS compatibility.

To establish the impact of CPA addition and dilution to cells in 96-well microtiter plate format, ten cell lines were selected for evaluation. Cells were added to the plates and incubated overnight to facilitate cell attachment. The following day, a solution of 4M CPA was added, followed by dilution 1 h later. Cells were imaged immediately before CPA loading, ∼30 min after CPA loading and immediately after CPA unloading. As shown in Figure 1, we observed significant cell detachment from two cell lines examined, namely LnCaP and HCT-8. The pattern of cell detachment is irregular, where in several lines large cell clusters containing both live and dead cells are released from the surface of the plate in a highly irregular manner (Supplemental Figure 1). This is not ideal as uncontrolled cell detachment is expected to result in well-to-well and plate-to-plate variability, which would reduce the robustness of the assay. Although other cell lines remained adhered after CPA challenge and dilution, seven were excluded from consideration due to practical limitations such as long doubling time, poor confluency, incompatibility with viability assays or sensitivity to CPAs (Supplemental Table 1). Ultimately a human bladder epithelial cell line T24, was selected based on its rapid doubling time, consistent confluency and stable attachment to the surface of the plate after recovery from CPA challenge. The osmotic resistance of T24 cell line [15] enables the addition of high concentrations of CPAs without cellular detachment, which is ideal for development of an HTS assay to assess CPA toxicity. Addition of CPAs in multiple steps increases assay time and may reduce throughput when scaling to test thousands of CPAs or CPA cocktails. While it is unclear why the T24 cell line is resistant to detachment, it may be due to its origin from bladder tissue, which experiences high osmotic gradients under normal physiological conditions. Further analysis of T24 osmotic sensitivity in the final assay format is described in section 3.6.

### 3.2 Assay Development: Establishing standard assay conditions for the T24 cell line

Several assay parameters were examined to establish the suitability of the T24 cell line for use in the CPA toxicity HTS assay. This includes identification of a robust endpoint assay, establishing the optimal cell seeding density, selection of the appropriate time point in which to assess survival following CPA exposure, selection of the carrier solution, and troubleshooting to reduce assay variability.

The MTT assay was utilized to assess cell survival following CPA exposure. As shown in Figure 2A, T24 cells were seeded at density of 4,000-40,000 cells/well with viability quantified next day using the MTT assay at estimated doubled density. These results confirm the cell density is within the linear range using MTT for quantification of viability. We also examined CellTiterGlo and alamarBlue, but neither assay provided robust results for assessing T24 viability following CPA exposure (Supplemental Figure 2).

Next, the appropriate endpoint for quantification of cell survival following CPA exposure was established. Here, GFP-labeled T24 cells were added to the 96-well plate followed by overnight incubation. The following day, 1,2-propanediol was added to cells at a concentration ranging from 0.5-5 M. The GFP signal was then quantified after 10min and 60min of CPA exposure. As shown in Figure 2B, no cell mortality is observed within this time frame. Following dilution of CPAs, some cell mortality is observed. Cells were then returned to standard cell culture conditions for overnight recovery. The following day, increased mortality is observed, indicating delayed-onset cell death, which has previously been widely reported following cryopreservation. [1,3] Based on this information, we selected an assay endpoint of 24h post-CPA removal. Of note, while useful for monitoring cell death in real time, the GFP- T24 cell line was not practical as an endpoint assay due to interference of Me_2_SO with signal acquisition.

Given the selected assay endpoint is 48h after cell seeding, we next established the optimal cell seeding density to ensure cells do not overgrow within this time range. We found an optimal cell seeding density of 20,000 cells/well (Supplemental Figure 3). This density ensures a monolayer of cells following overnight incubation, while remaining within the linear range of the assay signal 48 h later, when cell viability is quantified.

We further examined the effects of various vehicle carrier solutions in the presence of CPAs (Figure 2C, see also, Supplemental Figure 4). As shown for ethylene glycol, a carrier solution of DPBS- FBS-HEPES was found to maintain a physiologically relevant pH despite high concentrations of CPAs. Further incorporation of a centrifugation and aspiration step following CPA dilution led to more robust dose-response results when compared to the manual ‘flick-plate’ method that is standard for the MTT assay (Figure 2D).

### 3.3 Assay Validation: T24 CPA toxicity assay for HTS implementation

Prior to implementation, HTS assays are expected to undergo rigorous validation studies to ensure assay performance. [11] This is critical as well-to-well or plate-to-plate variability would interfere with assay reproducibility, potentially resulting in higher rates of false positives and false negatives. Therefore, in the HTS field, plate uniformity studies are typically conducted to derive statistical assay performance metrics, including the Z-factor [23], drift and CV. The Z-factor measures the statistical effect size of positive and negative controls, where a Z-factor >0.5 is considered an excellent assay due to the ability to reliably distinguish hits in large datasets based on their activity relative to the positive and negative controls. CV and drift are also applied to understand the assay variability among controls, where a CV and drift <20% is desired. Collectively, these metrics can validate assay quality and ensure reproducibility for the collection of high quality and informative data.

We examined the performance of the developed T24 CPA toxicity assay using statistical methods recommended by the National Institutes of Health Assay Guidance Manual for the development and implementation of HTS assays (assay logistics shown in Figure 3A). [11] Here, a checkerboard assay was performed (Figure 3B) [11,23] consisting of alternating control wells with a ‘low’ assay signal (5 M 1,2- propanediol, positive control) and a ‘high’ assay signal (no-CPA, negative control). By comparing the high and low assay signals, statistical measures can be applied to assess the quality and reliability of the assay and serves as an indicator of whether the signal window is large enough to distinguish hits from not-hits during HTS. Accordingly, a Z-factor of 0.75 was calculated for the checkerboard assay, well above the acceptable threshold of >0.5, indicating the assay is of ‘excellent’ quality. The assay was also examined for drift using standard methods. [11] Here, we found the maximal drift of the positive control was 7.5% and the negative control was 10.3%, where maximal drift was observed when plotting well number by column, then by row (Figure 3C). These results are well below the acceptable threshold of <20% drift. A favorable intra-assay CV was calculated at 5.9% for the negative (no-CPA controls), also below the acceptable threshold of <20%. The CV of the positive (CPA-treated) control was 22.5%, which is outside the <20% threshold expected for HTS assays. However, controls that exhibit minimal assay signal frequently fail the CV criterion [11], so the standard deviation (SD) is used as an alternative, where the guidelines state the SD of the low signal control should be less than that of the high signal control.

Accordingly, we found an SD of 0.03 for the positive (low signal) and 0.09 for the negative (high signal) controls, which meet the criteria for plate acceptability. Collectively, these data demonstrate the T24 CPA toxicity assay exhibits robust performance and is appropriate for HTS implementation.

### 3.4 Pilot Screen: Establishing dose-response toxicity of individual CPA solutions

Seven molecules previously used as CPAs or in CPA solutions were selected for further evaluation in the validated T24 CPA toxicity assay. A dose-response toxicity assay was performed where each CPA was tested at a concentration of 0.5-5 M in 0.5 M increments. The assay was repeated on three separate days where each CPA was tested in duplicate on each day to characterize reproducibility. The resulting LD_50_ curves (Figure 4) were then used to guide the formulation of CPA cocktails in the pilot screen. In our assay, toxicity followed the trend of urea > ME_2_SO > 1,2-propanediol > 3-methoxy-1,2- propanediol > glycerol > 1,3-propanediol > ethylene glycol. The LD_50_ of urea was ∼1.5 M, while ethylene glycol was only marginally toxic at even 5 M concentrations.

### 3.5 Pilot screen: Design and technical outcomes

Pilot screens (also called a mini-validation screens) are typically performed prior to the full HTS campaign (i.e. primary screen) to assess logistics. [5] The pilot screen consists of a smaller library and can identify barriers to assay implementation such as high hit rates or poor reproducibility. [5,17] This allows for the identification of issues under real-world screening conditions prior to execution of expensive and time-consuming primary screens. For our purposes, the pilot screen served to identify the optimal working concentration for CPAs in this assay, and to ensure that the selected assay conditions are appropriate for identification of hits. Therefore, we performed a pilot screen where CPA cocktails were tested at a range of concentrations to establish the dynamic range and hit rate, while characterizing reproducibility.

Using the validated HTS assay, a pilot screen was performed using mixtures of the seven CPAs. The plate layout is shown in Figure 5A, where each microtiter plate accommodates 42 experimental wells, with three columns of positive and negative controls and cell-free outer wells containing PBS to account for edge effects. The on-plate controls are used to calculate a Z-factor for each individual plate, where only plates that meet a Z-factor >0.5 are considered of acceptable quality. A signal window for identification of low toxicity CPAs was established at ≥80 % cell survival, where the assay achieves 91.6% sensitivity and 100% specificity. This threshold can be adjusted based on the desired balance between the false negative and false positive hit rate (Supplemental Figure 5). Due to the exploratory nature of the pilot screen, replicates were performed for each cocktail (N=2-9) to monitor overall assay variability between plates and over time. Further, replicate cocktails and single CPAs were occasionally included to monitor the effects of well location within the plate over time (Supplemental Figure 6). In total, 587 unique CPA cocktails were tested at concentrations ranging from 3.5-8 M (resulting in 2,352 total experiments completed), where each cocktail contained between two and seven components.

Out of 56 plates screened, 53 passed quality control (i.e. Z-factor >0.5) resulting in a 95% success rate. All three failed plates were screened on the same day, suggesting a technical issue that affected experiments conducted during that period. Among acceptable plates, the Z-factor ranged from 0.52-0.95, with an average of 0.77±0.10 and a median Z- factor of 0.78. The favorable outcome from the pilot screen combined with the validation study suggests this assay would be appropriate for primary HTS implementation.

A major goal of the pilot screen was to establish a hit rate at various CPA concentrations. Hits were identified here using a defined threshold ≥80% average cell survival. The highest hit rate of 63% was observed for the 3.5 M CPA cocktails. The lowest hit rate (0%) was observed among CPA cocktails at ≥6 M. The 5 M CPA cocktails exhibited a wide range of cell survival rates (i.e. 3-87%), with an overall hit rate of 1.7% (Figure 5C, Table 1). Overall outcomes and hit rates are tabulated in Figure 5E,F.

**Table 1.** Representative CPA cocktails examined at a concentration of 5 M. Of the 177 unique CPA cocktails tested at an overall concentration of 5 M, three hits were identified (noted with an asterisk), resulting in a 1.7% hit rate. The top ten performing CPA cocktails are highlighted in grey, based on highest rate of survival. The worst performing CPA cocktails, based on lowest survival rate, are shown in the lower half of the table. Of note, poorly performing cocktails containing urea were excluded from this analysis as this CPA was very toxic even at very low concentrations.

| CPA Cocktail | Glycerol | Me2SO | Ethylene Glycol | 1,2-propanediol | 1,3-propanediol | 3-methoxy-1,2-propanediol | Urea | Total molarity | Average % Survival | stdev |
| --- | --- | --- | --- | --- | --- | --- | --- | --- | --- | --- |
| 424* | 0 | 0 | 3.5 | 0.5 | 0.5 | 0.5 | 0 | 5 | 87.3 | 0.7 |
| 375* | 0 | 0 | 4.5 | 0 | 0 | 0 | 0.5 | 5 | 87.1 | 3.5 |
| 253* | 0 | 0 | 3 | 0 | 1 | 1 | 0 | 5 | 81.0 | 1.6 |
| 178 | 2 | 0.5 | 2 | 0.5 | 0 | 0 | 0 | 5 | 75.1 | 0.8 |
| 446 | 2 | 0.5 | 2 | 0 | 0.5 | 0 | 0 | 5 | 75.0 | 0.6 |
| 426 | 3 | 0.5 | 0.5 | 0 | 0.5 | 0.5 | 0 | 5 | 74.8 | 2.2 |
| 425 | 3 | 0.5 | 0.5 | 0.5 | 0.5 | 0 | 0 | 5 | 74.4 | 1.7 |
| 423 | 3.5 | 0 | 0 | 0.5 | 0.5 | 0.5 | 0 | 5 | 73.9 | 7.6 |
| 421 | 3.5 | 0.5 | 0.5 | 0.5 | 0 | 0 | 0 | 5 | 73.8 | 8.1 |
| 202 | 1 | 0.5 | 2 | 1.5 | 0 | 0 | 0 | 5 | 73.2 | 4.7 |
| 274 | 0.5 | 0 | 0 | 1.5 | 1.5 | 1.5 | 0 | 5 | 21.9 | 1.0 |
| 277 | 1 | 1 | 0 | 0 | 1.5 | 1.5 | 0 | 5 | 22.2 | 5.8 |
| 279 | 1 | 0 | 0 | 1 | 1.5 | 1.5 | 0 | 5 | 22.3 | 3.9 |
| 264 | 0 | 0 | 0 | 3 | 1 | 1 | 0 | 5 | 25.2 | 3.1 |
| 517 | 1.5 | 0.5 | 0 | 0.5 | 1 | 1.5 | 0 | 5 | 25.3 | 15.1 |
| 270 | 0.5 | 1.5 | 0 | 0 | 1.5 | 1.5 | 0 | 5 | 26.1 | 3.7 |
| 242 | 0.5 | 0 | 0.5 | 0.5 | 0.5 | 3 | 0 | 5 | 26.7 | 3.0 |
| 294 | 0 | 0 | 0 | 1.5 | 1.5 | 2 | 0 | 5 | 26.8 | 4.5 |
| 288 | 1.5 | 0 | 0 | 0 | 1.5 | 2 | 0 | 5 | 28.4 | 9.9 |
| 226 | 0 | 0 | 0 | 2 | 1 | 2 | 0 | 5 | 28.9 | 2.0 |

\*CPA cocktail identified as a hit (i.e. $\geq 80\%$ survival of T24 cells under assay conditions).

Figure 5B-D shows the results of CPA cocktails screened at an overall concentration of 4, 5 or 6 M (see Supplemental Excel file for detailed list of CPA cocktail compositions and associated viability values). Occasionally, data clustering is observed because of similarities in neighboring CPA composition. For example, in Figure 5D, the first eight CPA cocktails exhibit low survival compared to the last eight. This is due to ethylene glycol bias, as this is the lowest toxicity CPA screened based on our LD_50_ results (Figure 4). Specifically, the first eight CPA cocktails have very low concentrations of ethylene glycol in contrast to the final eight. Overall, the standard deviation of replicates was acceptable (averaging 5.4%), though we observed the highest standard deviation in plates that contained cocktails #127-168, which are included in Figure 5B. Here, the CPA cocktails were screened in duplicate (on separate plates) on the same day. As both plates had a passing Z-factor and no technical failures were noted during assay execution, the source of variability is unknown. However, of the 53 passing plates examined in the pilot screen, this was the only unexplained and systematic anomaly identified.

### 3.6 Pilot screen: Preliminary analysis of results

The goal of the pilot screen was to assess assay performance. However, several trends in toxicity were noted, and are briefly discussed below. To assist with this discussion, the predicted toxicity of each cocktail was calculated based on the dose-response toxicity for each individual CPA (Supplemental Excel File) for comparison to experimental outcome.

#### 3.5 M – 4.5 M CPA cocktails

Of the 293 CPA cocktails tested at lower concentrations (i.e., 3.5-4.5 M), cell viability ranged from 31-100%. The overall hit rate at this concentration range was ∼38%. While a high hit rate reflects low toxicity and is typically a positive outcome in the context of CPA toxicity, this presents a challenge for HTS, where the goal is to identify a manageable subset of hits for subsequent validation and optimization. While overall good agreement is observed among predicted and experimental toxicity, ∼177 CPA cocktails exhibited lower than expected toxicity. These cocktails most often contained some combination of glycerol, Me_2_SO, ethylene glycol and 1,2-propanediol.

#### 5 M CPA cocktails

Of the 177 CPA cocktails tested at 5 M, cell viability ranged from 3-87%. A favorable hit rate of 1.7% was observed (Fig. 5C). This hit rate is considered ideal as this would lead to a more manageable set of hits in the primary screen, where many thousands of cocktails may be screened. Further, given the dynamic range of cell viabilities, the 5 M concentration may provide an informative dataset for the purpose of identifying patterns and relationships between CPA compositions and their biological activities, including approaches through data mining and machine learning. While 5 M CPAs may not be sufficient for vitrification of large volumes, such as organs, ≤5M solutions have been used to vitrify a range of cell types. [6,9,12,19] Table 1 lists the composition of ten 5 M CPA cocktails that exhibit the lowest toxicity and ten that exhibit the highest toxicity. On average, these 177 CPA cocktails are more toxic than predicted. To ensure toxicity was not due to osmotic damage, we randomly picked several 5 M CPA cocktails that were low toxicity, moderate toxicity, or high toxicity. These were rescreened using a 5-step CPA loading and 5-step CPA unloading procedure. We observed strong agreement between the single- and multi-step results (Supplemental Figure 7), suggesting that cell mortality is related to CPA toxicity rather than osmotic sensitivity. Collectively, these data demonstrate that screening cocktails at 5 M concentration can yield a robust data set with appropriate dynamic range, reproducibility, optimal hit rate and absence of osmotic damage. Cocktails that that exhibited lower than expected toxicity were more likely to contain a mixture of glycerol, Me_2_SO, ethylene glycol and 1,2-propanediol. Cocktails that were more toxic than predicted were more likely to contain urea. However, no firm conclusions are made based on this pilot screen due to the small sample size. These trends in toxicity will be further explored in the future primary HTS screen.

#### ≥6 M CPA cocktails

At 6-8 M, 111 CPA cocktails were examined, but no hits were identified (see Fig. 5D for 6 M cocktails). All but two of the cocktails were more toxic than predicted. It is unclear whether this is due to osmotic stress, or non-specific CPA toxicity, which occurs when excessive hydrogen bonding with water may lead to dehydration of macromolecules. [8] While no hits were identified at this concentration, it is expected that toxicity could be reduced through incorporation of shorter durations of CPA exposure, multi-step CPA loading, or lower temperatures. While we anticipate the assay can be readily modified for compatibility with these assay conditions, it was avoided here as a reduction in toxicity would lead to a high hit rate or the need for multi-step CPA loading, which is not desirable as our goal is scale up to a full HTS campaign. However, these conditions will be further explored in the future lead identification and optimization stages of the HTS process (Schematic 1).

## 4. Conclusions

Here, we report the development of an HTS compatible assay for the characterization of CPA toxicity, where the first three steps of the HTS approach were completed including assay development, validation, and completion of a pilot screen. These steps are typical in the HTS field prior to execution of the primary screen to minimize false-positives and false-negatives and to assess assay performance and logistics. The developed assay includes T24 cell monolayers in 96-well format with on-plate controls which allow for assessment of plate performance. Successful validation was demonstrated using statistical metrics developed for the HTS field including Z-factor, CV, and drift. [4,11] A pilot screen of 587 unique CPA cocktails (N=2-9 replicates; 2,352 total experiments) was then performed to understand well-to-well and plate-to-plate variability of the assay. Among the 56 plates tested, a 95% success rate was achieved based on an acceptable Z-factor of >0.5. We found excellent reproducibility across plates and on different days. Further analysis of assay results indicates that screening at the 5 M cocktail concentration yields a high dynamic range (3-87% viability) and manageable hit rate (1.7%). Further, we found that the 5 M CPA cocktails can be added in a single step with no observation of osmotic sensitivity. While multi-step CPA loading is practical when screening a small number of microtiter plates, it becomes limiting when the assay is utilized for the primary HTS campaign, where we intend to manage upwards of 20 plates per day.

In the future, we intend to complete a larger HTS campaign to provide a robust toxicity dataset that may be useful for data mining, machine learning, and identification of hits for further optimization. Hits will be characterized in several ways. This includes the execution of secondary assays, where hits will be tested against additional cell lines, and characterization of glass forming tendencies. These secondary assays will require modifications in assay execution, including for example, multi-step CPA loading and unloading. However, since the number of hits is expected to be much smaller than the number of cocktails tested in the primary screen, more complicated assay logistics are acceptable. Secondary screens are expected to enable the prioritization of the most promising CPA cocktails for further characterization. We further anticipate this assay may be useful to screen for new potential CPAs.

## Data statement

The datasets generated during and/or analyzed during the current study are available on request from the corresponding author.

Selection of appropriate cell line, experimental conditions, positive and negative controls, assay readout, etc.

Determine whether assay is suitable for HTS using prevailing statistical methods (e.g. drift, CV, Z-factor, plate-to-plate and well-to-well variability, etc.)

Perform mini-validation screen to assess assay performance (e.g. reproducibility, Z-factor, hit rate, dynamic range, etc.) across broad assay conditions.

Perform large-scale screening campaign (≥thousands of CPAs or CPA cocktails).

Cherry-pick hits to confirm activity; optionally- perform dose-response analysis.

Assess performance of hits in additional biospecimens (e.g. cell lines, tissues, etc.) to identify those with broad activity (optional).

Hits are optimized based on temperature of exposure, stepwise loading/unloading, length of CPA exposure, etc. Cocktails can be optimized in dose-response manner if desired.

