## Supplemental Tables and Figures for "Validation of a high-throughput screening assay for the characterization of cryoprotective agent toxicity"

**SUPPLEMENTAL INFORMATION**

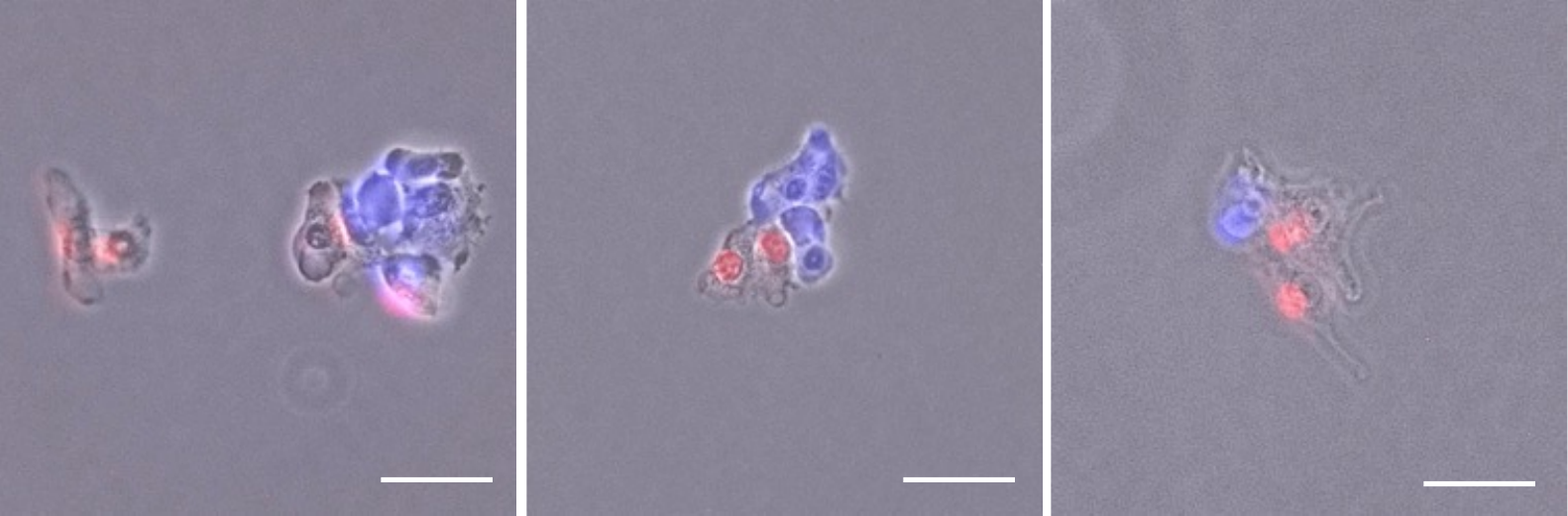

**Supplemental Figure 1**. **Clusters of LnCaP cells detached from the microtiter plate during CPA dilution.** Clusters contain both dead (PI^+^, red) and live (Calcein AM^+^, blue) cells. Scale is 50 µm.

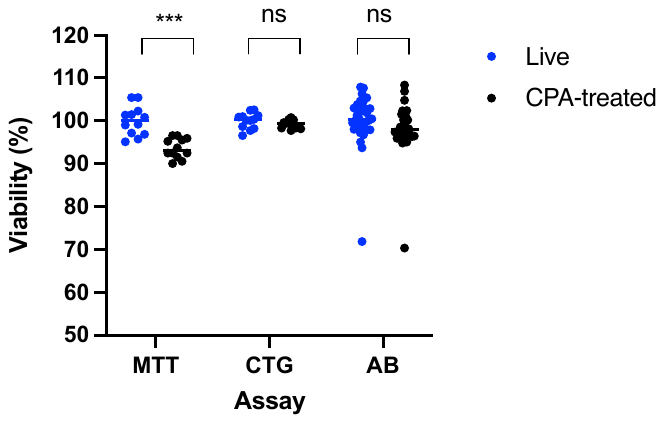

**Supplemental Figure 2. Comparison of viability assays.** The MTT, CellTiter-Glo (CTG) and Alamar Blue (AB) viability assays were examined for compatibility with the CPA toxicity HTS assay. Briefly, cells were plated onto a 96-well microtiter plate as described in the methods section. The viability of cells with no CPA (negative control) was compared to cells treated with a low-toxicity CPA cocktail consisting of 1.5 M glycerol and 2 M ethylene glycol. The MTT assay successfully distinguished between live and CPA-treated T24 cells (t-test, p < 0.0001). However, the CTG and AB assays were not sensitive enough to detect significant differences between control and CPA-treated wells (t-test, p = 0.21 and p = 0.81, respectively).

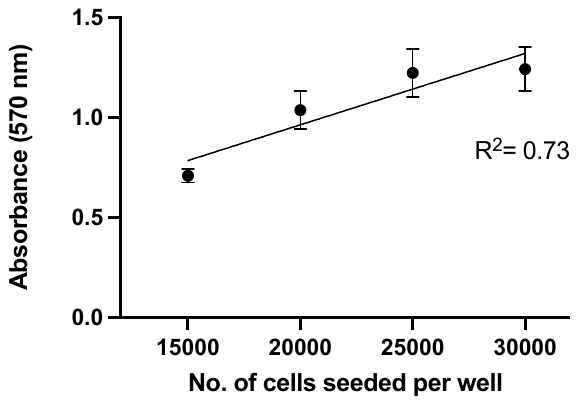

**Supplemental Figure 3. T24 standard curve measured 48 h after seeding by the MTT viability assay.** T24 cells were seeded at increasing densities and assayed after 48 hours, corresponding to the CPA toxicity HTS assay endpoint, by which time the cell density was estimated to have quadrupled. The observed cell density remains within the linear range (R^2^ =0.73).

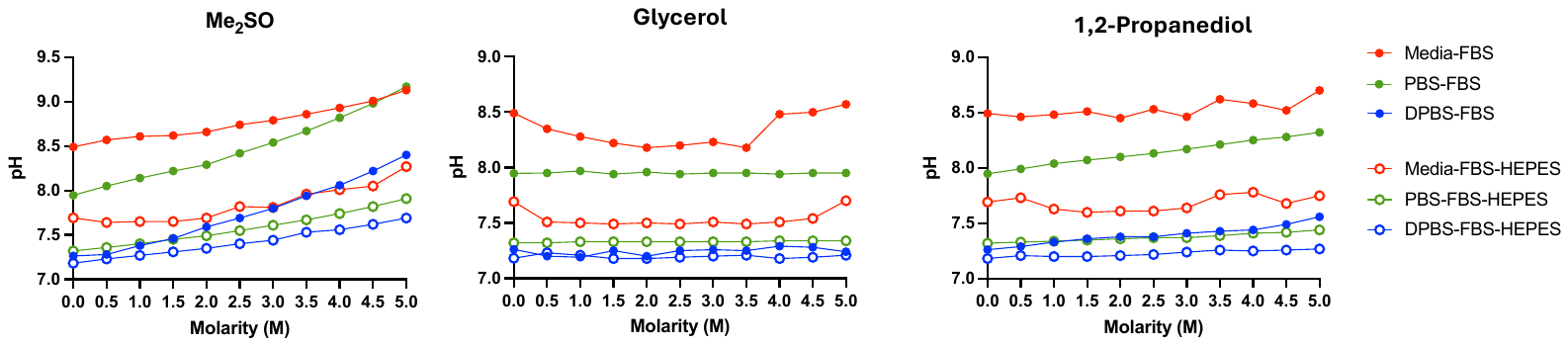

**Supplemental Figure 4. Examination of the impact of vehicle carrier solutions on pH of CPA cocktails.** DPBS supplemented with HEPES maintains pH of ME_2_SO, glycerol and 1,2-propanediol cocktails across concentrations. Curves for ethylene glycol is shown in Figure 2C of the main text.

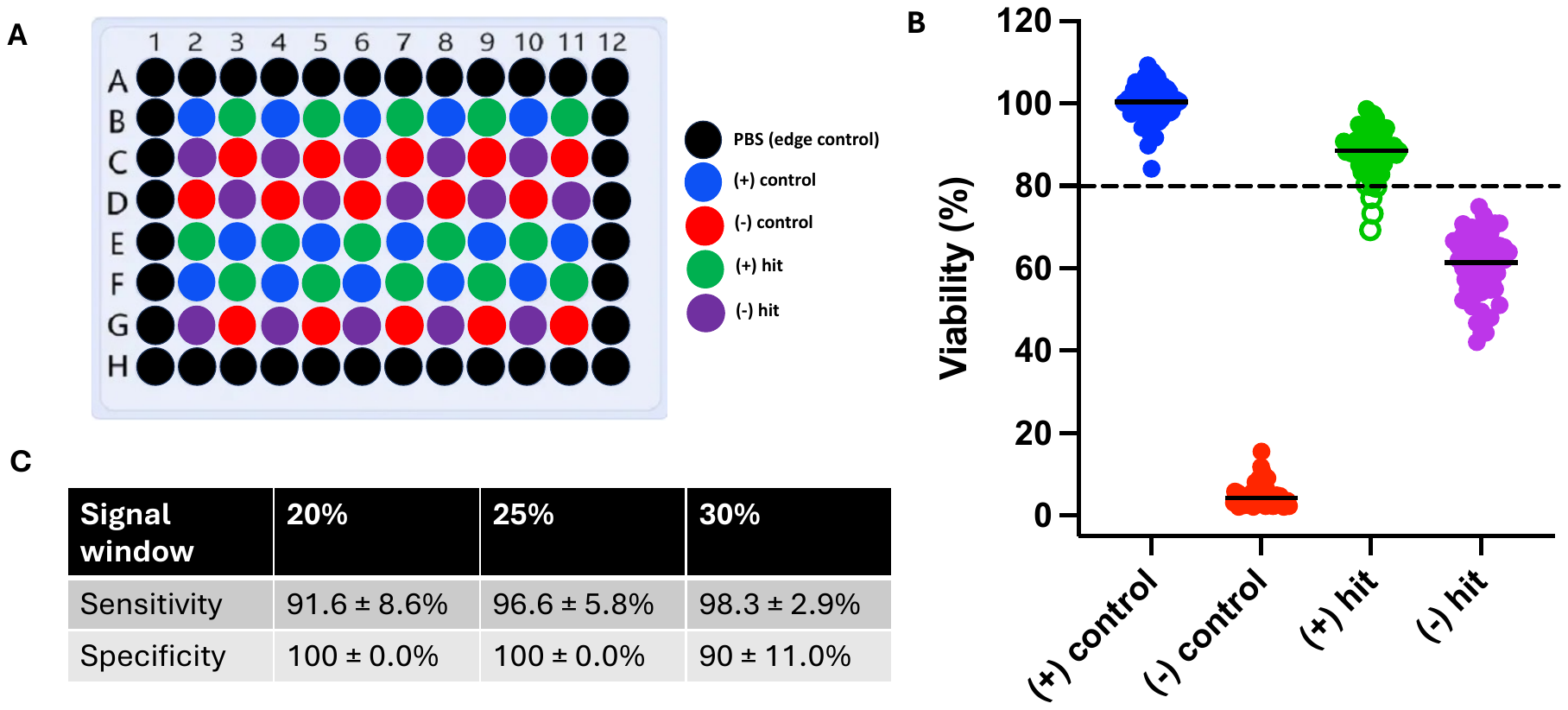

**Supplemental Figure 5. Impact of viability threshold on CPA toxicity HTS assay performance metrics. (A)** A checkerboard assay was designed consisting of alternating wells containing T24 cells: live, (No CPA, (+) control), dead (5 M 1,2-propanediol, (-) control), treated with a known low toxicity CPA (1.5 M glycerol/ 2 M ethylene glycol, (+) hit) and treated with a known moderate toxicity CPA (1.5 M glycerol/ 2 M ME_2_SO, (-) hit). Outer wells were unseeded and contained PBS to account for edge effect. **(B)** Cell viability threshold enables identification of positive hits (above the threshold) and negative hits (below the threshold). **(C)** Adjustment to cell viability threshold influences the desired balance between false negative and false positive hit rates, as evidenced by variations in assay sensitivity and specificity. Sensitivity refers to assay ability to accurately identify true positive hits, while specificity pertains to the accurate identification of true negative hits.

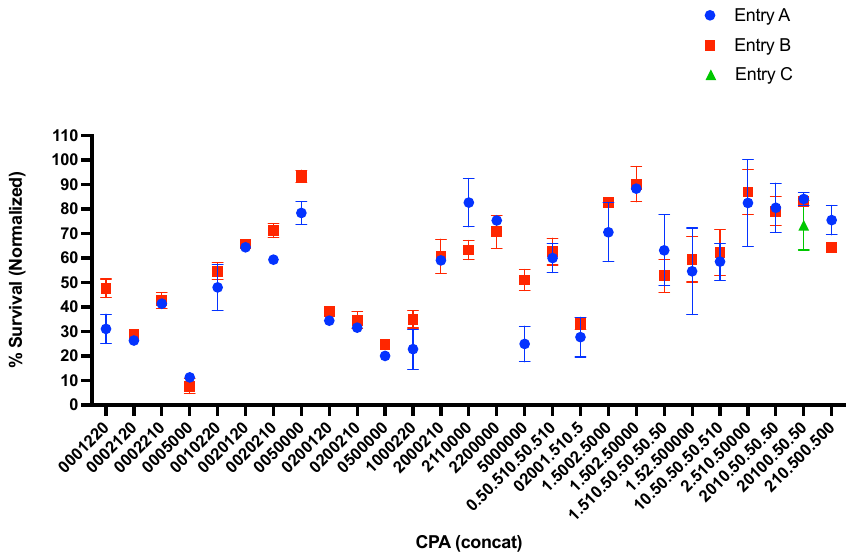

**Supplemental Figure 6. Comparison of CPA cocktail duplicates.** Single CPAs and CPA cocktail duplicates were included at random intervals throughout the pilot screen to measure the effects of well placement on T24 cell survival. Cell survival is plotted as the average and standard deviation as a function of the CPA identity. Overall, good agreement among duplicate CPA cocktails is observed.

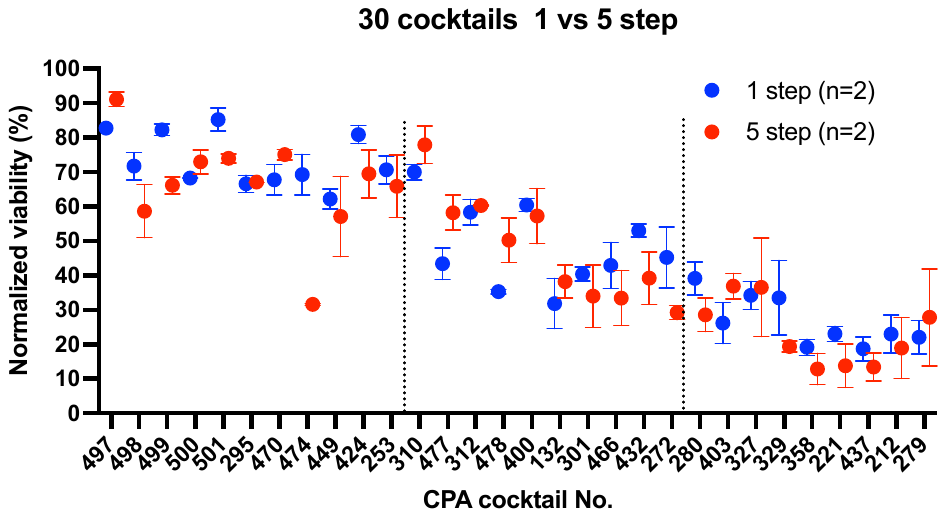

**Supplemental Figure 7. Comparison of single- and multi-step CPA loading.** Following completion of the pilot screen, we selected a cohort of CPA cocktails that exhibit low, moderate, or high toxicity for secondary analysis based on the effects of CPA stepwise loading. Here, CPAs were compared on the same day (N=2) where loading was accomplished in either a single step, or in five step-wise additions. CPA unloading was then accomplished in either two steps (for single step CPA loading), or in five steps (for multistep CPA loading). CPA cocktails ranged from 4-5 M in each category of low, moderate or high toxicity. See Supplemental Excel File for cocktail compositions.

**Supplemental Table 1. Summary of ten cell lines screened for compatibility with the CPA toxicity HTS assay.**

| **Cell type** | **Species** | **Tissue type** | **Doubling time** | **Reason for exclusion** |
| --- | --- | --- | --- | --- |
| **HEPA 1-6** | Mouse | Hepatocellular carcinoma | 27 h | Doesn`t reach 100% confluence |
| **LnCaP** | Human | Prostate carcinoma | 60 h | Doesn`t reach 100% confluence  Sloughing off |
| **HCT-8** | Human | Colorectal adenocarcinoma | 28 h | Sloughing off |
| **PC3** | Human | Prostate adenocarcinoma | 25 h | Resistant to MTT permeabilization |
| **NCI-H1650** | Human | Bronchoalveolar carcinoma | 42 h | Long doubling time |
| **SK-BR-3** | Human | Breast adenocarcinoma | 38 h | Long doubling time  Doesn`t reach 100% confluence |
| **HUVEC** | Human | Primary umbilical vein endothelial cells | 36 h | Long doubling time  Senescence  Sensitivity to CPA |
| **MSC** | Human | Mesenchymal stem cells | 26 h | Sloughing off  Loss of phenotype after 4-5 passages and differentiation into myofibroblasts |
| **MDBK** | Bovine | Kidney epithelium | 35 h | Long doubling time  Sensitivity to 1,2-propanediol |
| **T24** | Human | Bladder carcinoma | 19 h | -- |
